# A causal inference framework for estimating genetic variance and pleiotropy from GWAS summary data

**DOI:** 10.1101/531673

**Authors:** Yongjin Park, Liang He, Manolis Kellis

## Abstract

**Motivation:** Much of research in genome-wide association studies has only searched for significantly associated signals without explicitly removing unwanted source of variation. Confounder correction is a necessary step to reveal causal effects, but often skipped in a summary-based analysis.

**Results:** We present a novel causal inference algorithm that controls unwanted sources in genetic variance and covariance estimation tasks. We demonstrate substantially improved statistical power and accuracy in extensive simulations. In real-world applications on the UK biobank summary statistics data, our method recapitulates well-known pleiotropic modules, suggesting new insights into biobank-scale GWAS analysis.

**Contact:** YP and MK

## I. INTRODUCTION

### a. Motivation

In the advent of nation-wide biobank databases, genome-wide association studies (GWAS) have been conducted on unprecedentedly diverse and massive sets of complex traits. For instance, in the UK biobank [41], thousands of traits are associated with genetic variants, and the summary statistics data are made publicly available for follow-up studies. Leveraging this massive information is important for us to understand biological mechanisms of human biology, and eventually, to achieve precision and personalized medicine.

Of many important properties, *polygenicity* and *pleiotropy* are repeatedly and commonly observed in the past decades of GWAS data analysis [43], compelling that these two ideas should be properly modeled. Under the polygenic inheritance model, in stark contrast to traditional monogenic regime, a large number of genes (variants) drive variability of downstream phenotypes; even though a single individual variant may exert only a small fraction of total effect, an aggregate effect accounts for most of genetic variance. It is not difficult to realize that a univariate (SNP-by-SNP) GWAS test is fundamentally limited in statistical power and interpretability, but common practice of GWAS, even phenome-wide association studies (PheWAS), has not completely moved out of the univariate paradigm.

Pleiotropy is pervasive across multiple types of human traits, it is no longer expected to have a single GWAS variant fully committed to a single trait. For instance, genetic variants near *PCSK9* gene are widely associated with many different human traits, including lipid metabolism, cardiovascular disorders, type 2 diabetes, and Alzheimer’s disease [10]. Pleiotropic patterns can emerge for many reasons. Underlying regulatory and metabolic pathways are commonly perturbed by genetic variants. Or, we may simply observe them because definitions of human traits are redundant and elusive. Yet, by knowing genetic underpinnings of pleiotropic patterns, we can improve our predictions of potential adverse and beneficial side effects of drugs, and even refine definitions of human traits.

Calculating genetic variance and genetic covariance between traits is perhaps a foremost important step in multi-trait GWAS analysis as they directly measure polygenicity and pleiotropy, respectively. By locally calculating them, we improve our resolution. We establish a set of causally associated traits in comparison with many related traits in biobank, and uncover novel comorbidity networks with clear conviction of relevant genomic locations.

### b. Problem definition

We focus on estimating these second-order statistics from summary data. Existing summary-based methods, agnostic to data generation process, are unable to characterize and adjust biases introduced by non-genetic effects. Most methods inevitably depend on hard-coded assumptions and only address special cases of confoundedness [13, 32, 38, 39, 45]. However, we are concerned that a substantial proportion of estimated genetic variability may contain contributions from non-genetic effects, such as cryptic relatedness [6]. Especially, for cross-trait analysis on a single cohort, such as UK Biobank, where samples are inevitably shared, we are even more concerned that traits are easily confounded by uncharacterized effects.

## II. APPROACH

In this work, we present a novel causal inference method, `RUV-z` (Removing Unwanted Variation in GWAS z-score matrix), with which we characterize undesired sources of information lurking in summary statistics, and selectively remove them to improve accuracy and statistical power of local variance/covariance calculation.

We first introduce `zQTL` (z-score based quantitative trait locus analysis), a suite of machine learning (ML) methods for summary-based regression and matrix factorization, then demonstrate how we can successively apply the factorization and regression steps to design a new confounder-correction method (Alg1).

Our approach is inspired by existing methods, established on individual-level data analysis in genomics [11, 36] and astrophysics [37]. We conceived our approach asking, “What could have been done if we had fully observed information?” and carried over the core concepts into summary-based data analysis. Nonetheless, to our knowledge, `RUV-z` is the very first attempt for explicit confounder correction in summary statistics data analysis in GWAS.

## III. METHODS

### A. Backgrounds

#### a. Generative model for individual-level phenotypes

We model a quantitative trait of *n* individuals were generated by a multivariate linear regression model on *n* × *p* genotype matrix *X* measured on *p* common genetic variants:

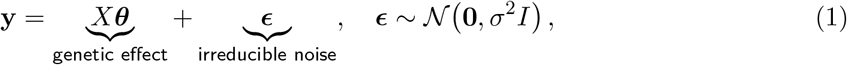

where we assume non-genetic variance follows the isotropic Gaussian distribution with unit residual variance *σ*^2^. For a case-control study, we may consider **y** as a liability score. And we only observe summary GWAS statistics effect size and standard error for each *j* ∊ [*p*]:

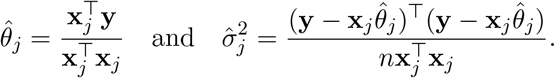

From reference cohort, we can easily estimate the LD matrix, *R* = *n*^−1^*X*^⊤^*X*, using column-wise standardized genotype matrix *X*.

#### b. Generative model of GWAS summary statistics

For simplicity, letting 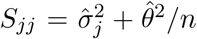, we can redefine the model with respect to *p*-dimensional summary statistics-the regression with summary statistics (RSS) model [45]:

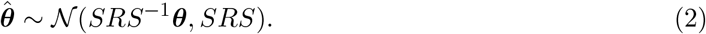

Normally we have large enough sample size (*n* → *∞*), the RSS model resorts to a fine-mapping model [16]. Generative scheme of a GWAS z-score vector, with each element 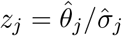, is described by the reference LD matrix *R* and true (multivariate) effect size vector *θ*.

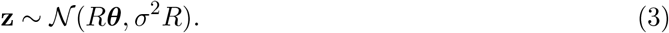

##### 1. Variance calculation

*a. Local heritability estimation* Two notable works have been introduced to calculate local heritability from GWAS summary statistics-(1) spectral approach [38] and (2) Bayesian approach [45]. Both method rely on the same multivariate Gaussian recession model, but differ in terms of statistical inference algorithm.

*b. Spectral approach* The method suggested by [38] uses a regularized infinitesimal polygenic model, assuming all the variants weakly contribute to polygenicity, and therefore all SNPs are included in the model. Suggested estimation of multivariate polygenic vector is simply 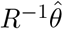, yet in a typical situation, estimated LD matrix 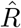 is rank-deficient. [38] suggests using pseudo-inverse matrix (or regularized matrix) from singular value decomposition of reference genotype matrix. With the results of singular value decomposition, *n*^−1/2^*X* = *UDV*^τ^, the inverse matrix can be approximated, *R*^−1^ = *VD*^−2^*V*^τ^, at a certain rank *q*, and this yields the following heritability estimation:

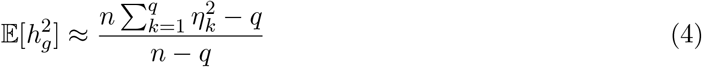

where *n* is sample size of the underlying GWAS and 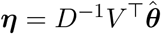.

*c. Bayesian approach* On the other hand, [45] takes a more direct approach to carry out posterior inference of the multivariate effect sizes *θ*, treating GWAS summary statistics as data of the RSS model (Eq.2). Once the posterior distribution of *θ* become available, we can easily characterize the total variance of polygenic effects,

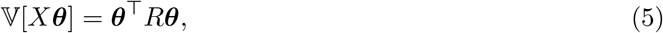

over the runs of Markov chain Monte Carlo steps. However, Bayesian approaches typically scale poorly especially when we pose sparse prior over the effect sizes, such as “spike-slab” [27] to select a relevant set of causal variables probabilistically. We need to explore over *O*(2^*p*^) combinatorial space.

##### 2. Covariance calculation

For a pair of standardized trait vectors, such as ***y_s_*, *y_t_***, we estimate covariance between trait t and s by 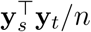. Expanding the statistics with respect to the multivariate model (Eq.1),

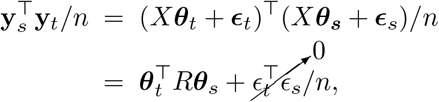

we may derive a closed form solution of the genetic covariance in summary statistics [25].

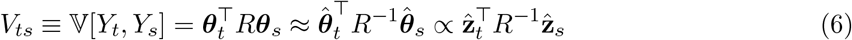

where the last approximation stems from a polygenic infinitesimal model estimated from the GWAS summary vectors, i.e., 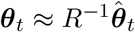 and 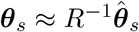. Since the *R* matrix can be rank-deficient, we use the same technique used in [39]. Overall, we found the rank *q* = 50 worked well in most cases, simulated by reference panels of European ancestry.

### B. Model-based characterization of unwanted effects

#### 1. Definitions

We modify the original definition of phenotype model (Eq.1), by introducing an additional term u, which accounts for overall effects of confounding variables (Fig.1a, c). For each trait *t* of total *r* traits, we have

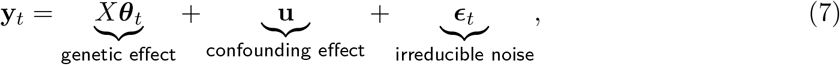

where we assume there is a single confounding vector **u** and independent errors *∊*, and the distribution follows

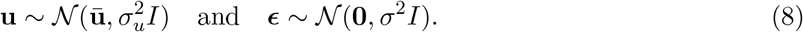

**FIG. 1:**
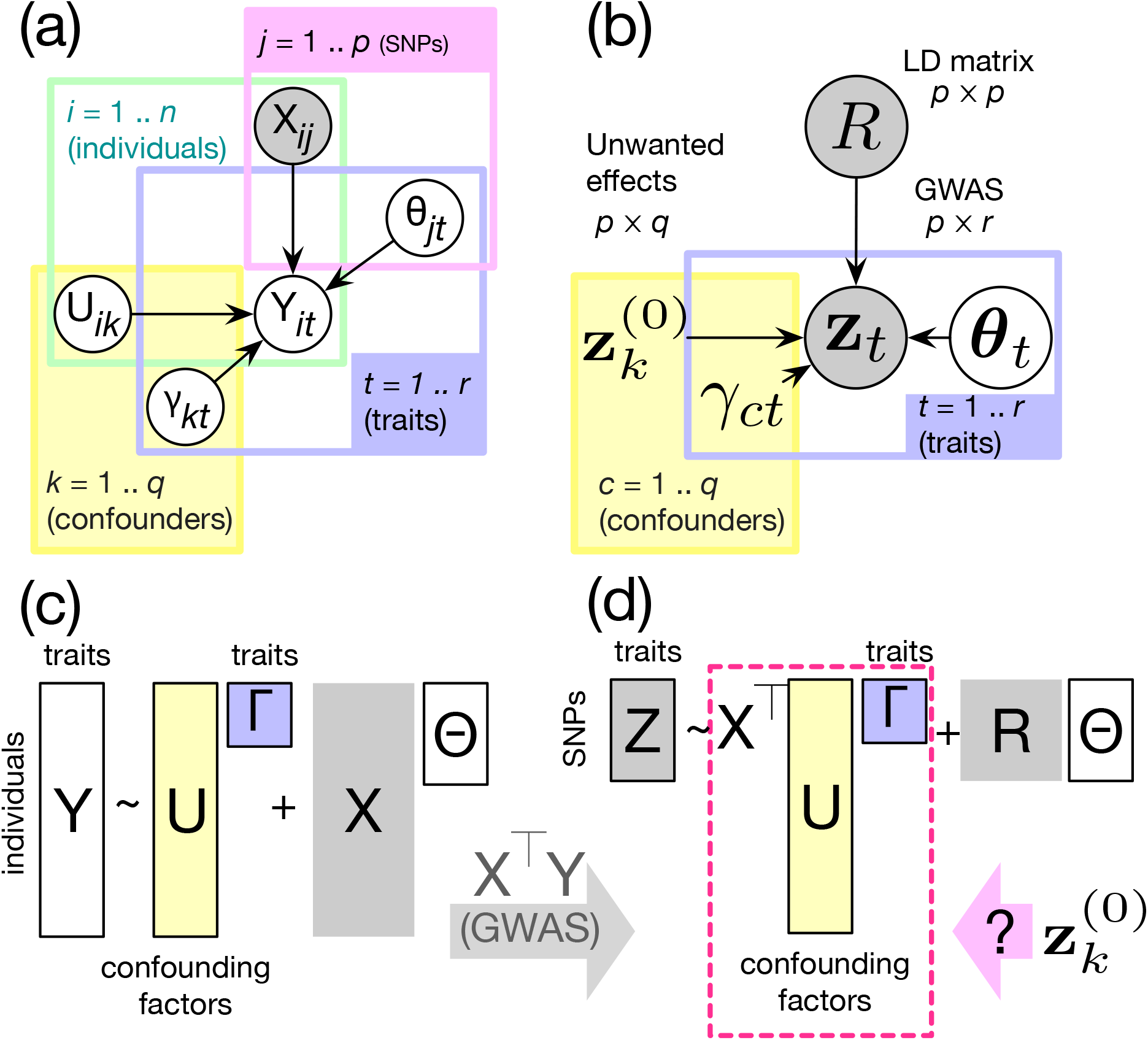
Illustration of multi-trait generative models for individual-level phenotype (a, c) and GWAS summary statistics data (b, d). *n* number of individuals; *p*: number of SNPs; *q*: number of confounding factors; *r*: number of traits; *X*: genotype matrix, *n* × *p*; *U*: confounding covariates, *n* × *q*; *Y*: unobserved phenotype matrix, *n×r*; *θ* sparse genetic effect size, *p* × *r*; *γ*: loading matrix for confounding factors, *q* × *r*. **(a)** A graphical model for individual-level data generation, **(b)** Summary-based formulation for the graphical model (a), **(c)** A matrix view of the model (a), **(d)** A matrix view of the model (b); this model can be derived from (c) by multiplying genotype matrix *X*^⊤^. Our primary concern is to estimate legitimate surrogate z-scores 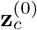 to account for the effects of confounding effects *X*^⊤^*U*.

Just like the previous z-score model (Eq.3), we can easily derive the distribution including the confounding factor terms (Fig.1b, d).

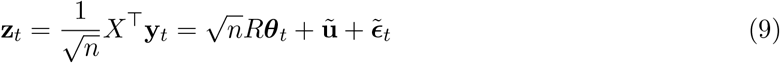

where the second (confounding effect on the z-scores) and third variables (multivariate error covariance by LD structure) follow:

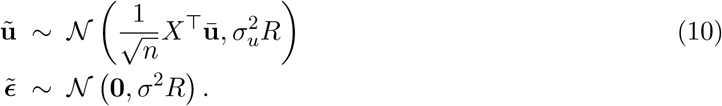

In this work, we explore two possible forms of this *u* variable: (1) polygenic bias and (2) non-genetic confounding factors shared across traits.

#### 2. When do we worry about confounding effects?

##### a. Polygenic bias by weak directional pleiotropy

We address a special type of the confounding effect modestly correlated with genetic information, as a tenant of the column space of *X*, which can introduce unwanted directional pleiotropy. Following the definition of [2], the mean parameter *ū* of Eq.10 takes a special form, **ū** = *ūX*^τ^**1**_*p*_/*p* with *ū* ≠ 0. We broadly term this type of effects “polygenic bias” because we would have

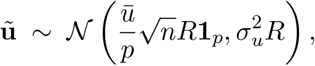

and this result in biased estimation of the multivariate effects in a polygenic fashion,

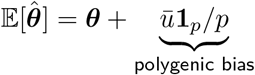

in any model estimation.

##### b. Non-genetic confounding variables shared across traits

Unlike the trait-by-trait heritability estimation, unwanted bias can be introduced in the covariance estimation (Eq.6), even if we had unbiasedness of the confounding variable, 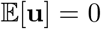. We can expose such a case algebraically. Under this confounding effect model (Eq.7), covariance estimate between trait *t* and *s* can be rewritten with respect to the model parameters:

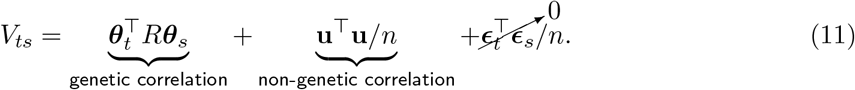

Since 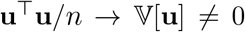, although 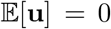, we would get fundamentally biased covariance estimation.

In practice, we never have full access to the individual-level confounding effects, **u**, but we can trace back the results of confounding effects observed in GWAS z-score matrix. We define surrogate z-score vectors 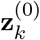 to capture unwanted correlations 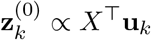. In the the z-score model (Eq.3), we can include these confounder terms, **z**^(0)^:

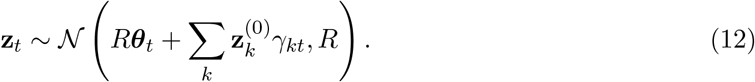

where we impose Bayesian sparsity [27] on the multivariate effect size parameters, meaning that *a priori* 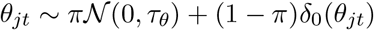 and 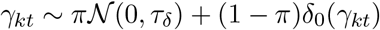.

#### 3. How do we synthesize the surrogate z-scores to control bias

##### a. Characterization of polygenic bias

We address polygenic bias easily by including two “intercept” terms in the above regression model (Eq.12):

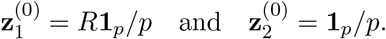

In Mendelian Randomization, similar ideas have been proposed (e.g., [2]). It also is straightforward to include other types of bias factors induced by genomic features such as minor allele frequency, GC content bias, and localized population structures, and so on.

##### b. Low-rank matrix factorization to identify a general class of confounding factors

Since hidden confounding effects of z-score originate from individual-level variations (Fig.1d), as illustrated in the previous derivations (Eq.9 and 10), in this matrix factorization, we seek to characterize and decompose latent factors on *hypothetical* individual-level phenotype matrix *Y*. Provided that *n* × *m* phenotype matrix 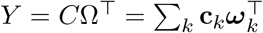, our goal is to find the confounding factors *C* (*n* × *q*) with the corresponding loading matrix Ω (*r* × *q*) at some rank *q* as small as possible. For each column *t* (trait) of GWAS z-score matrix, we simply formulate this factorization as:

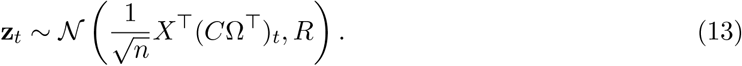

Enforcing Bayesian group sparsity [15] on each factor *k* ∊ [*r*], 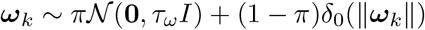, we approximate low-rank factorization in a data-driven way such that a selected non-zero factor pervasively affects on a majority of traits.

Although we can hardly expect the inferred latent factors align with actual individual-level confounding variables, they can be used to identify inter-trait covariance structures induced by non-causal confounding effects, such as **z**^(0)^ α *n*^−1/2^**X**^⊤^**c**_*k*_ for some factor *k*.

However, this latent factor may become correlated with genuine genetic effects. In the model (Eq.12), we can hardly expect that ***Rθ*** is perfectly orthogonal to some inferred confounding factor *n*^−1/2^**X**^⊤^**c**_*k*_, even after we adjust *R*. Here, we avoid such a possibility by carefully constructing a “control” data matrix 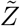, on which we can warrant orthogonality with a genotype matrix, and use this for matrix factorization. In terms of the traditional RUV methods [11], we may consider this 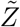, as “control gene” data matrix.

##### c. Causal inference to identify non-genetic confounding effects

To establish a legitimate control data 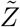, we make two causal assumptions:

1. *Independence across different LD blocks*: We expect genetic effects from different LD blocks are independent. For two genotype matrices *X*_1_ and *X*_2_ sampled from different LD blocks, we have 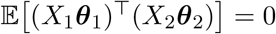 for any multivariate effects ***θ***_1_ and ***θ***_2_.
2. *Genomic location invariance of non-genetic effects*: On the other hand, non-genetic effects consistently exist without genetic association.

Suppose we construct a control data for some LD block *l*, taking advantage of an independent LD block *k*. We collect two z-score matrices, *Z_l_* and *Z_k_*, and two reference genotype matrices, *X_l_* and *X_k_*, respectively. Then, we get a proxy z-score matrix by this operation.

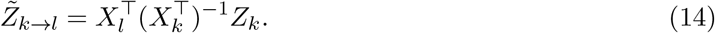

In practice, the matrix inversion is ill-defined, so we also use the truncated SVD technique [38], but we make use of overall spectrum of singular values up to some numerical stability (> 10^−3^). We carry out factorization (Eq.13) with this matrix 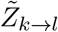 to profile confounding effects on the LD block *l*, i.e., 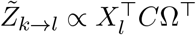.

#### 4. Model-based variance and covariance estimation

##### a. Variance (heritability) estimation

From the posterior inference results (Eq.12), it is straightforward to estimate un-scaled version of variance on each component.

- The genetic component: 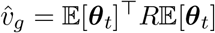;
- the covariate component: 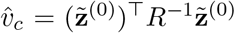, weighting contribution of each confounder differently, 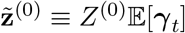;
- the residual component: 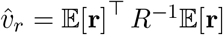 where we obtain posterior mean of the residual in estimation of the model, 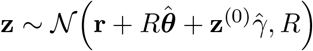, fixing other components.

From these, we estimate local heritability,

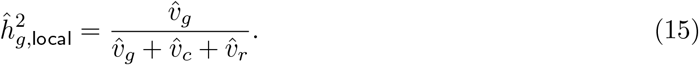

Our estimate is derived from the posterior inference results, so one can estimate variance by hundreds of parametric bootstrap steps. Explicit variance decomposition results (Eq.15) easily extend to genome-wide heritability estimation. Combining each LD block *l*’s variance estimates, namely, 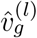, 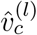, and 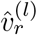, we calibrate genome-wide heritability estimation by the ratio of the overall quantities:

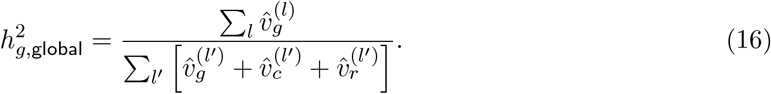

##### b. Covariance calculation

After the Bayesian inference of the sparse effect sizes on trait *t* and *s*, we obtain posterior mean and covariance to construct test statistic by the dot-product between the two effect sizes. Under the normality, we define the distribution of this dot-product between traits *t* and *s* as follows.

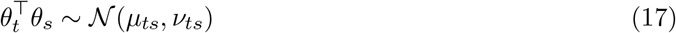

with analytical derivation [4]:

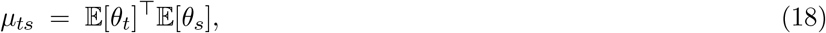

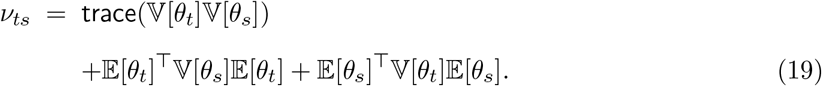

In the Wald test, we evaluate significance of local pleiotropy by rejecting the null hypothesis, 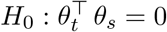 between traits *t* and *s*.

#### 5. Overall algorithm

Alg.1 summarizes overall steps in summary-based mult-trait analysis pipeline. As for the posterior inference of regression and factorization models, we use a modified version of stochastic variational inference algorithm [30]. See details in the supplementary text.

### C. Simulations

We simulate realistic phenotype matrix Y using imputed genotype matrix. We repeat full experiments on randomly selected 100 LD blocks, predefined by [3]. The details of simulation steps are outlined in Alg.2.

In simulation (Alg.2), we define genetic pleiotropy rather strictly that two traits are considered pleiotropic if and only if they share common genetic mechanisms in the SNP-level resolution.

## IV. RESULTS

### A. Simulations

#### a. RUV-z shows superior statistical power in causal trait discovery

In each set, we simulated summary data of total *r* = 100 traits, and seeded only 10 of them are causally associated, using

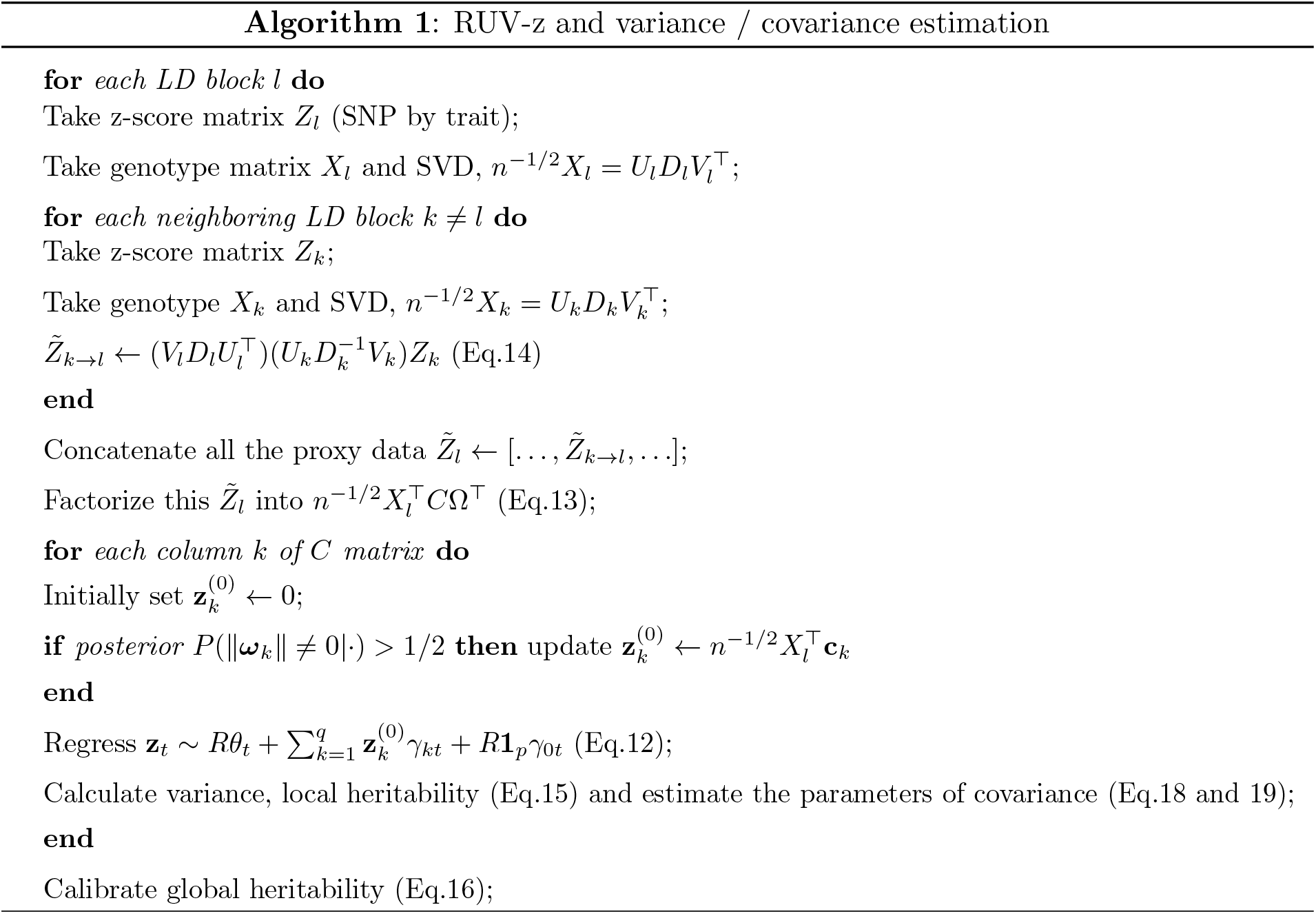

UK10K reference panel with n=6,285 [19]. We generate polygenic bias **u** as previously suggested [2] and feed into the simulation algorithm (Alg.2). We first randomly select 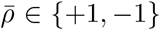 with probability 1/2, then fix the **u** vector by setting **u** ← *X_**ρ**_* with each element sampled, 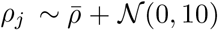. Based on the summary data, we compare accuracy of the following methods to estimate local heritability and causal trait discovery.

- The spectral method (Eq.4), applying different levels of SVD truncation (*q* = 10,50) and no truncation at all (full).
- The RUV-z method (Alg.1) with model-based estimation (Eq.15).

Here, we only show a representative simulation result, generated by 3 causal SNPs with 30% of variance induced by the polygenic bias component (Fig.2), but more comprehensive simulations results can be found in the supplementary material.

**FIG. 2:**
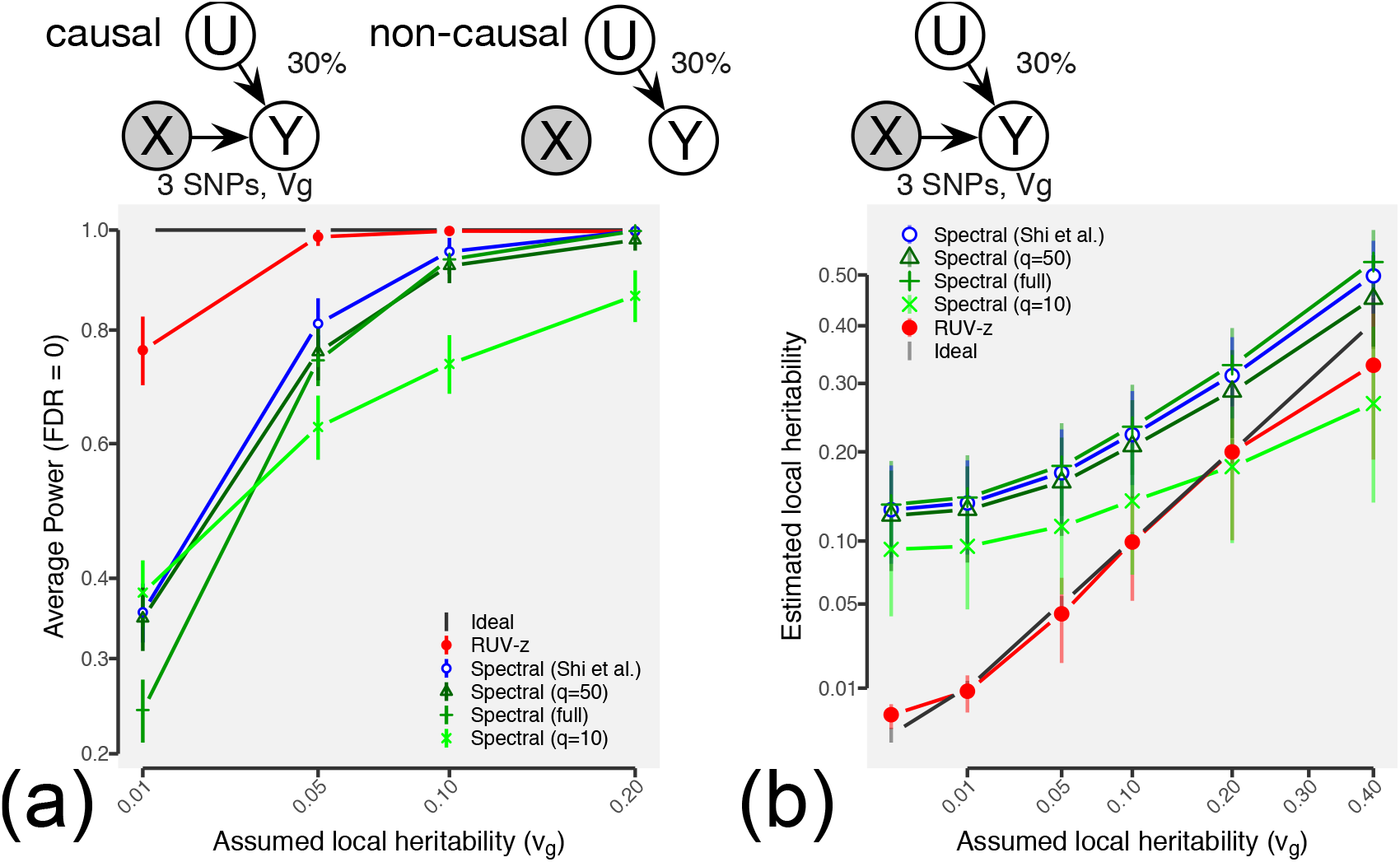
Model-based heritability calculation accurately prioritizes causally associated traits. We simulated data with three randomly selected SNPs varying different level of heritability (x-axis) under the influence of moderate polygenic bias (30% of total variance). *Shapes and colors*: different heritability estimation methods; *error bars*: 2 times of standard errors. **(a)** Power comparison. **(b)** Heritability estimation.

Across all the simulation settings, we find our RUV-z method outperforms the existing method. We may interpret our results in terms of model selection problem of underlying polygenic model. We think sparse modeling of the RUV-z method clearly separate strong genetic effects apart from polygenic bias component prevalently present across genome. Even at the lowest level of local heritability (0.01), our RUV-z method achieves nearly 80% power with no mistake. For the spectral methods, both under- and over-fitting (q=10 and full) damages statistical power, but the original spectral method [38] adaptively truncates the rank to yield marginally superior statistical power.

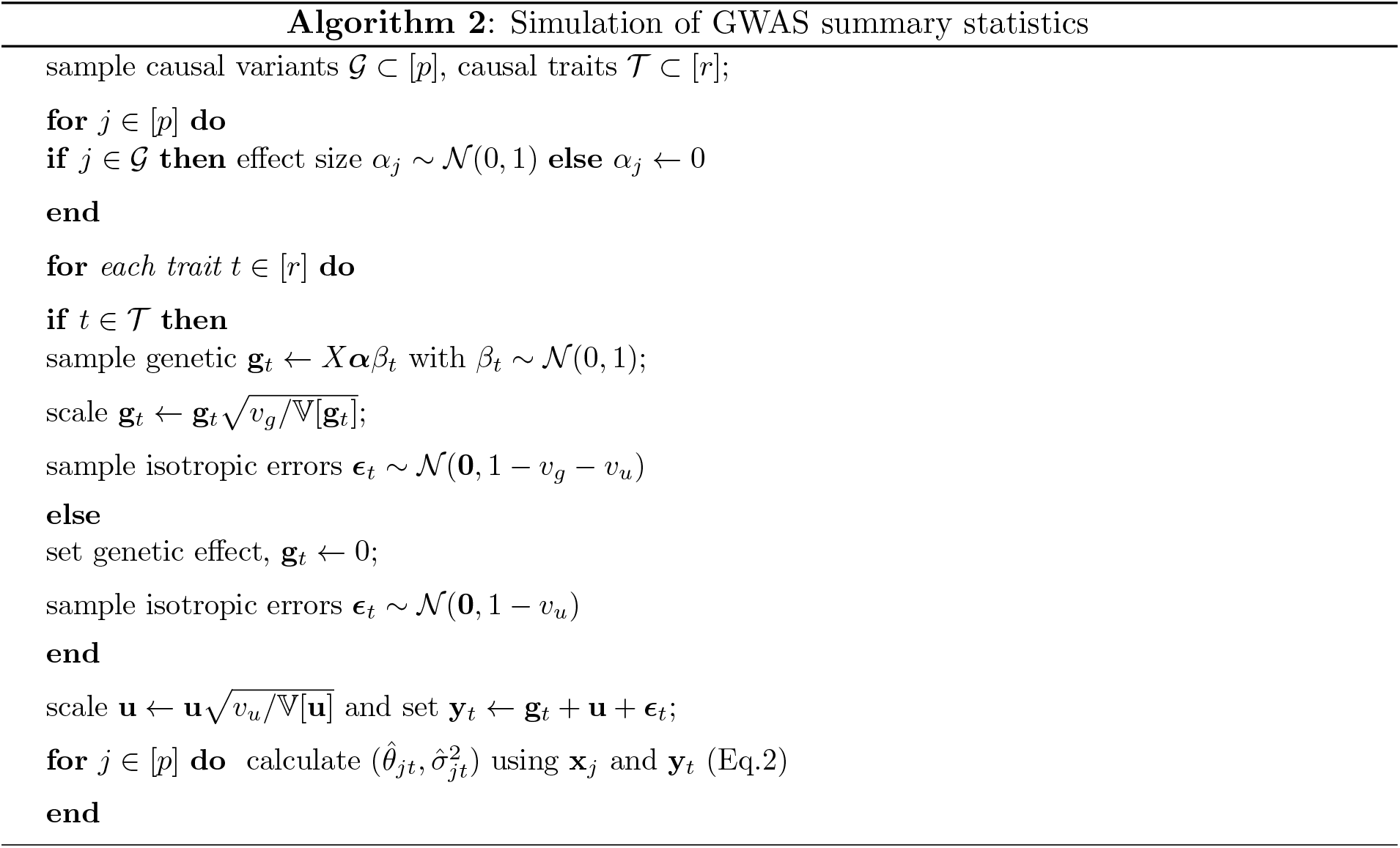

#### b. RUV-z achieves accurate heritability estimation without inflation

From the heritability estimation results, we can further emphasize that finding rightful complexity within a flexible framework is crucially important in polygenic modeling. We only find our Bayesian method can properly calibrate actual level of local heritability against the spurious polygenic bias (Fig.2b; see the supplementary results). Unlike the Bayesian method selects a relevant set of causal variants with the spike-slab prior [27], apart from polygenic bias terms **z**^(0)^, the spectral method is incapable of enforcing SNP-level sparsity, and its underlying model is fundamentally no different from a multivariate Gaussian model. Applying an aggressive level of SVD truncation may alleviate the inflated estimation due to the polygenic bias (q=10 in Fig.2b), but this rather flattens the overall trajectory of heritability estimation, elevating false discovery rates.

#### c. Sparse covariance estimation by RUV-z method outperforms other methods

In the covariance estimation problem, we specifically investigate each method’s capability to stand against the influence of non-genetic factors. We use a rather easy confounding effect **u**, sampled from isotropic Gaussian distribution 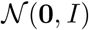. We use the same simulation (Alg.2), but this same confounding effect is shared used across all the 30 traits. Of them, 10 are causal traits; thus, of the total 435 trait-trait pairs, only 45 (10%) are truly pleiotropic. The confounding effects **u** account for 50% of variance.

On each simulated data, we compare three types of trait-trait scores:

- Raw: We calculate the summary-based genetic covariance matrix (Eq.6) without any adjustment, fixing the SVD truncation *q* = 50 or no truncation.
- Corrected: We first estimate potential non-genetic confounding effects (Alg.1), and regress them out from the raw z-score matrix, 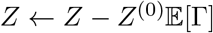, and then calculate the same type of genetic covariance on the adjusted z-scores (Eq.6), fixing the SVD truncation *q* = 50 or no truncation.
- RUV-z: We also estimate more direct statistics between the multivariate effect sizes (Eq.18).

In Fig.3(a) and (b), we present power comparison conducted on two reference panel genotype matrix, UK10K [19] and the 1000 genomes cohort (1KG) with European ancestry [42]. We have sufficiently large sample size (n=6,285) for the UK10K data, we have much fewer samples (n=502) on the 1KG data, therefore underpowered. On the UK10K data, even at the low level of local heritability (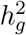 = .01), the covariance estimations with proper SVD-truncation (*q* = 50) easily achieve nearly 25-50% power without making a mistake (Fig.3a), whereas only 0-25% power can be seen in the 1KG data (Fig.3b).

**FIG. 3:**
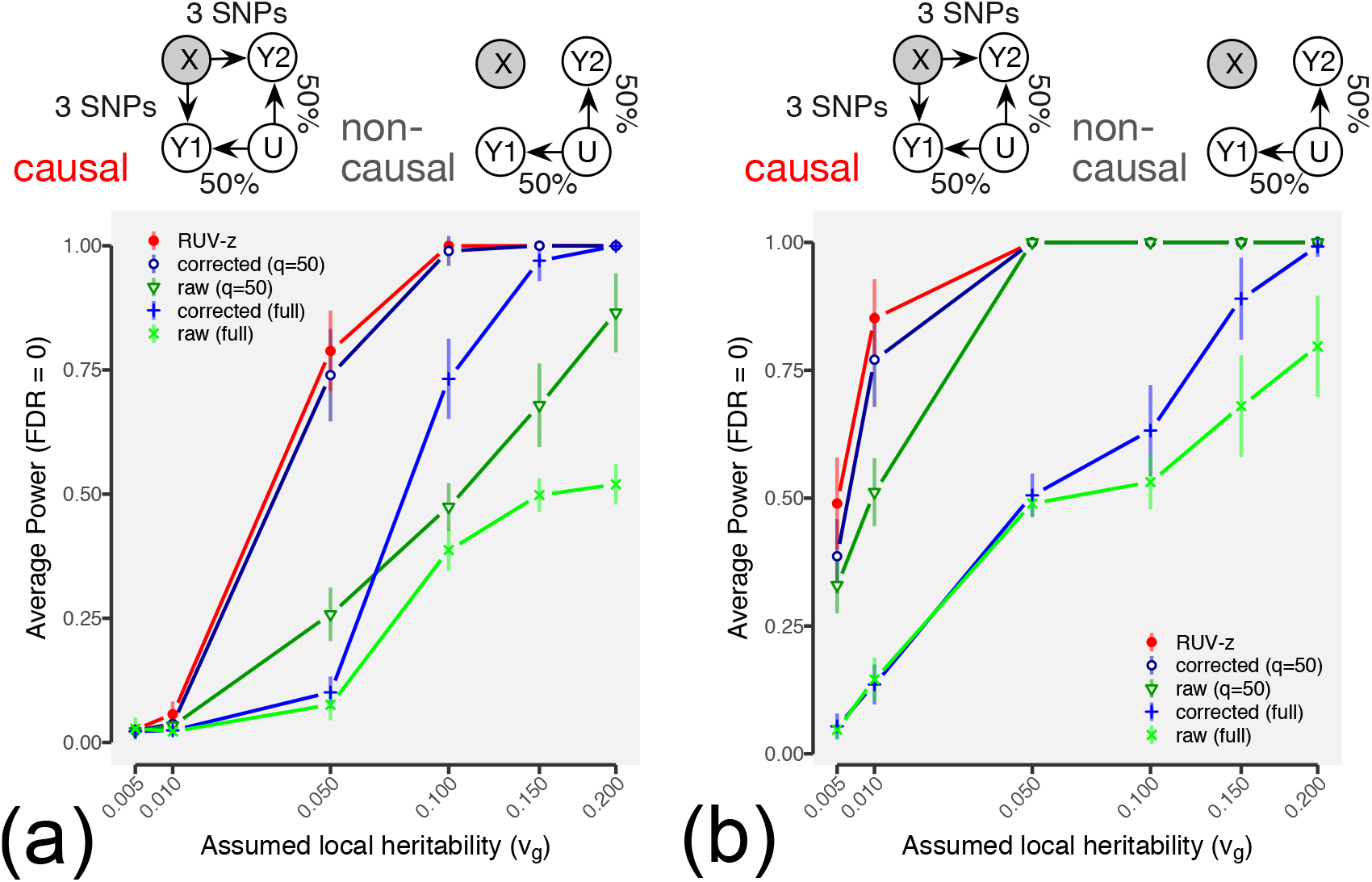
Summary-based local covariance estimation can be biased by the existence of unchracterized non-genetic confounding variables. We show the power comparison (y-axis) on the simulated data with 3 randomly selected SNPs at different level of heritability (x-axis) under the influence of moderate non-genetic effects (50% of variance). **(a)** Simulation using UK10K cohort [19] and **(b)** the 1000 genomes with European ancestry [42]. *Shapes and colors*: different heritability estimation methods; *error bars*: 2 times of standard errors (of 100 repetitions).

**FIG. 4:**
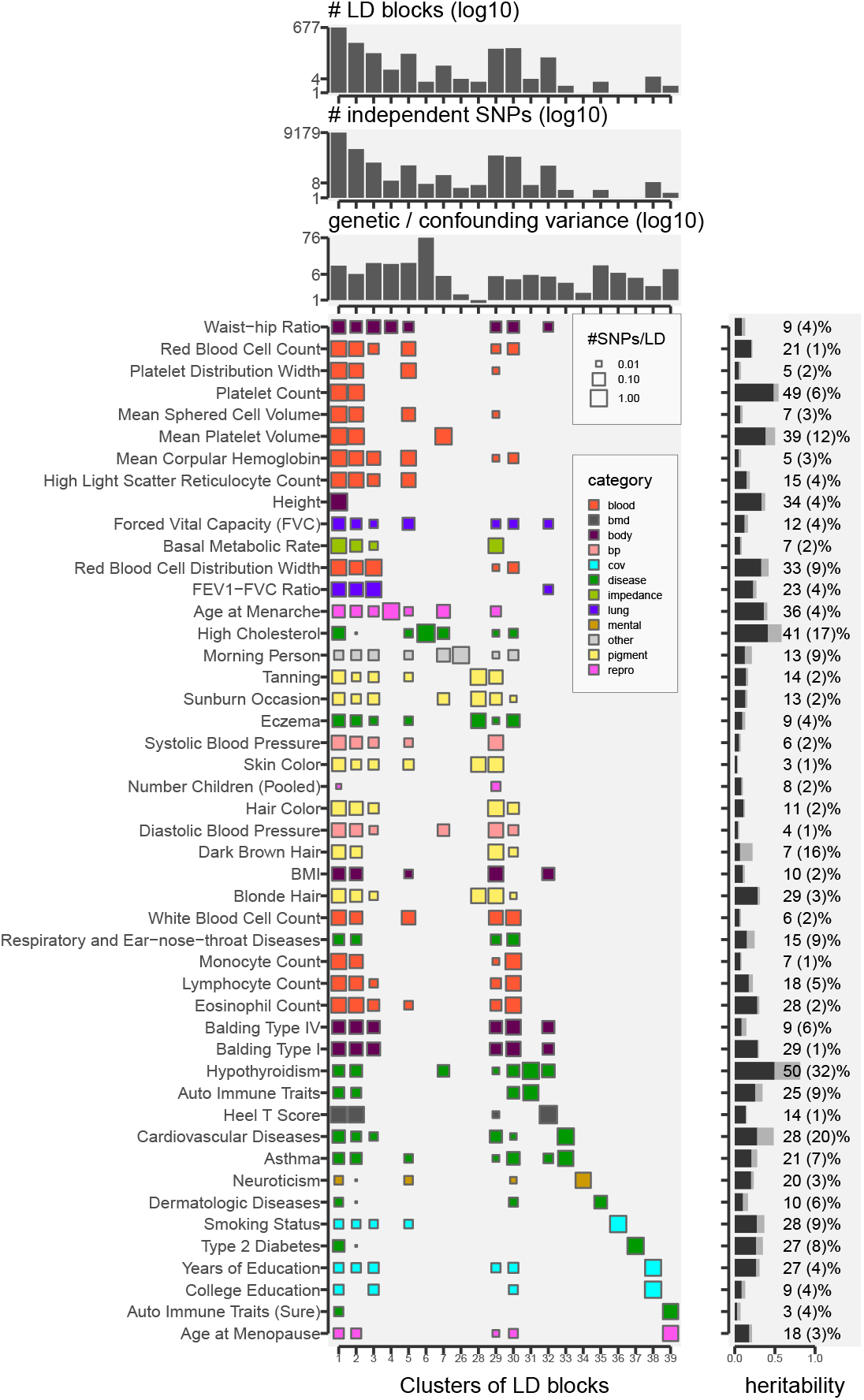
Clustering analysis of the LD blocks show the polygenic architecture of 80,041 trait-LD block pairs (of 47 UK Biobank traits and 1,703 LD blocks). Top three panels, from the 1st to the 3rd barplots, respectively show the sizes of each cluster in terms of number of LD blocks, number of statistically independent SNPs, and average ratio between the genetic and confouding effect variance. To better visualize low values, we scaled the y-axes in log10. The Hinton diagram (the 4th panel from the top) shows average number of causal SNPs within each cluster (column) on each trait (row) where the sizes of dots are scaled accordingly, and colored differently according to the broader category of traits previously used in the GWAS data [24]. On the right panel, each trait is annotated by the total genome-wide heritability (the dark gray bar with the percentage) and non-genetic confounding effects (the light gray bar with the number in the bracket).

Substantial impacts of uncharacterized non-genetic confounders can be seen in all the cases. Especially on the 1KG results at moderate local heritability (5%), the difference between the corrected and uncorrected method can exceed beyond 50% power; under the high heritability regime (> 5%), the confounder-corrected covariance calculation can achieve much higher power than any of the uncorrected calculations tuned with the level of SVD truncations. We can also confirm the sparse covariance estimation (Eq.17) after the RUV-z consistently outperforms other spectral methods (Eq.6). The sparse covariance, however, is designed to test more strictly defined pleiotropy than the conventional genetic variance, and its power depends on the accuracy of variable selection steps of the multivariate model estimation (Eq.7, or 9).

### B. Case study of UK Biobank (UKBB) traits

We investigated real-world GWAS summary statistics data, measured on the 47 common and complex phenotypes in the UKBB cohort [24] suing BOLT-LMM method [23]. We applied the RUV-z pipeline (Alg.1) on each of the 1,703 LD blocks [3] with the LD matrix estimated from the UK10K reference panel [19]. We locally calculate heritability, trait-trait covariance and number of causal SNPs across multiple traits. On each LD block, to learn hidden confounding effects, 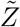, we used the information of neighboring 4 blocks (2 from the left; 2 from the right). Yet, we found final heritability estimations are largely invariant to the choice of the different number of neighbors that we tested 10 and 20.

#### a. Common genetic variants explain average 18% of phenotypic variance

From the total heritability estimation (Eq.16), we find a large fraction of phenotypic variances are explained by sparse multivariate effects of the common genetic variants, ranging between 3 and 50% with median 14.23% and mean 18.17% (± SD 12.32%). Our method based on sparse multivariate models appear to under-estimate total heritability of the anthropomorphic traits, such as height (34%), hair color (11%) and BMI (10%), compared to the original LMM-based results [24]. We may blame the lack of statistical power on our Bayesian inference, but we also think some proportion of discrepancy can be explained by latent covariates. Polygenic bias and non-genetic trait-trait confounders account for 1 - 32% of total variance with median = 3.95% and mean = 5.65% (± SD 5.81%).

Interestingly, nearly 50% of the total hypothyrodism variability can be explained by sparse genetic effects, but 32% of the variance can be also explained by potential confounding effects. Several twin studies [8, 14, 31] confirms that heritability of hypothyrodism (concentration of thyroid stimulating hormones) can reach over 60%, and it is also well accepted hypothyrodism is an outcome of other autoimmune disorders [8].

#### b. Clustering analysis reveals pleiotropy and comorbidity

We consider each LD block having a causal effect on a particular trait if and only if it contains any variant with the posterior inclusion probability (PIP) of the multivariate effect vector *θ* (Eq.12) exceeds 1/2. We obtained total 80,041 unique LD block-trait pairs and constructed sparse a feature matrix *W* (LD blocks × traits). For each element takes *W_it_* = 1 if there is a causal SNP on trait *t*, but *W_it_* = 0 otherwise. We resolved clustering of the 129 LD blocks with at least one causal trait association. Between the pairs of LD blocks (rows) we measure similarity by Jaccard coefficients; for instance, between a pair *i* and *j*, we calculate the similarity 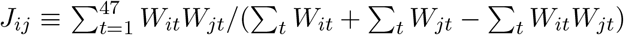.

We identified 20 non-empty clusters of LD blocks, and these LD blocks are not uniformly assigned to the 20 clusters (average size = 128.64 ± SD 117), rather the size distribution was seen highly skewed toward heavily pleiotropic groups. The cluster #1 spans over 677 LD blocks, involving 9,179 causal SNPs (this number can be loose; see the discussion).

The blood traits are genetically decomposed into multiple groups in relations with the other traits (the clusters #1, #2, #3, #5, #7, #29, #30). The clusters #5 and #30 clearly contrast with each other: The cluster #5 identifies genomic regions associated with the blood cell traits related to respiration and the forced vital capacity, whereas the cluster #30 combines the immune-related blood cells with the relevant common diseases.

However, we refrain from making a premature conclusion that these traits in the same group are regulated by shared biological networks, but only suggest this type of pervasive pleiotropic patterns observed in a large genomic region, and should be carefully dissected at a SNP-level resolution [12, 13, 17]. In fact, in our preliminary analysis (Park *et al.,* in preparation), these giant groups can be decomposed into multiple independent components by summary-based factored regression analysis [34].

#### c. Sparse multivariate covariance estimation uncovers nearly 8K pairs of local pleiotropy

Before we test local pleiotropy of trait pairs across LD blocks, following [39], we restricted our analysis on the LD blocks where both traits are strongly associated, therefore the selected LD blocks should explain substantial fractions of phenotypic variances on both traits. To be consistent with our clustering analysis, we called a trait is causally associated with a certain LD block with respect to the Bayesian variable selection of causal variants (PIP > 1/2).

Of the total 1,840,943 possible pairs of traits across 1,703 LD blocks, only 7,765 trait pairs (0.4%) are significantly correlated with respect to the multivariate effect sizes (Eq.17) at FDR < 5%. Of them, 4,228 pairs are positively correlated; 3,537 are negatively correlated (Fig.6). Sparse pleiotropy analysis may seem to yield overly conservative results, leading to considerably smaller number of discoveries than that 14,820 significant pairs (0.8%) are found by the summary-based covariance test (SVD truncation q=50) without any adjustment (Eq.6).

We claim that genetic covariance estimation without any adjustment, even with strong SVD truncation, can be seriously confounded by non-genetic covariates, because we can observe the following two things on the average covariance structure between traits (Fig.5). First, two types of average covariance matrices-one from the marginal and the other solely on the confounding effects-are highly similar to each other (Fig.5a and b). Here, we fix the same order of rows and columns for better illustration. But the covariance structure induced from the sparse effects (Fig.5c) exhibits no obvious similarity.

**FIG. 5:**
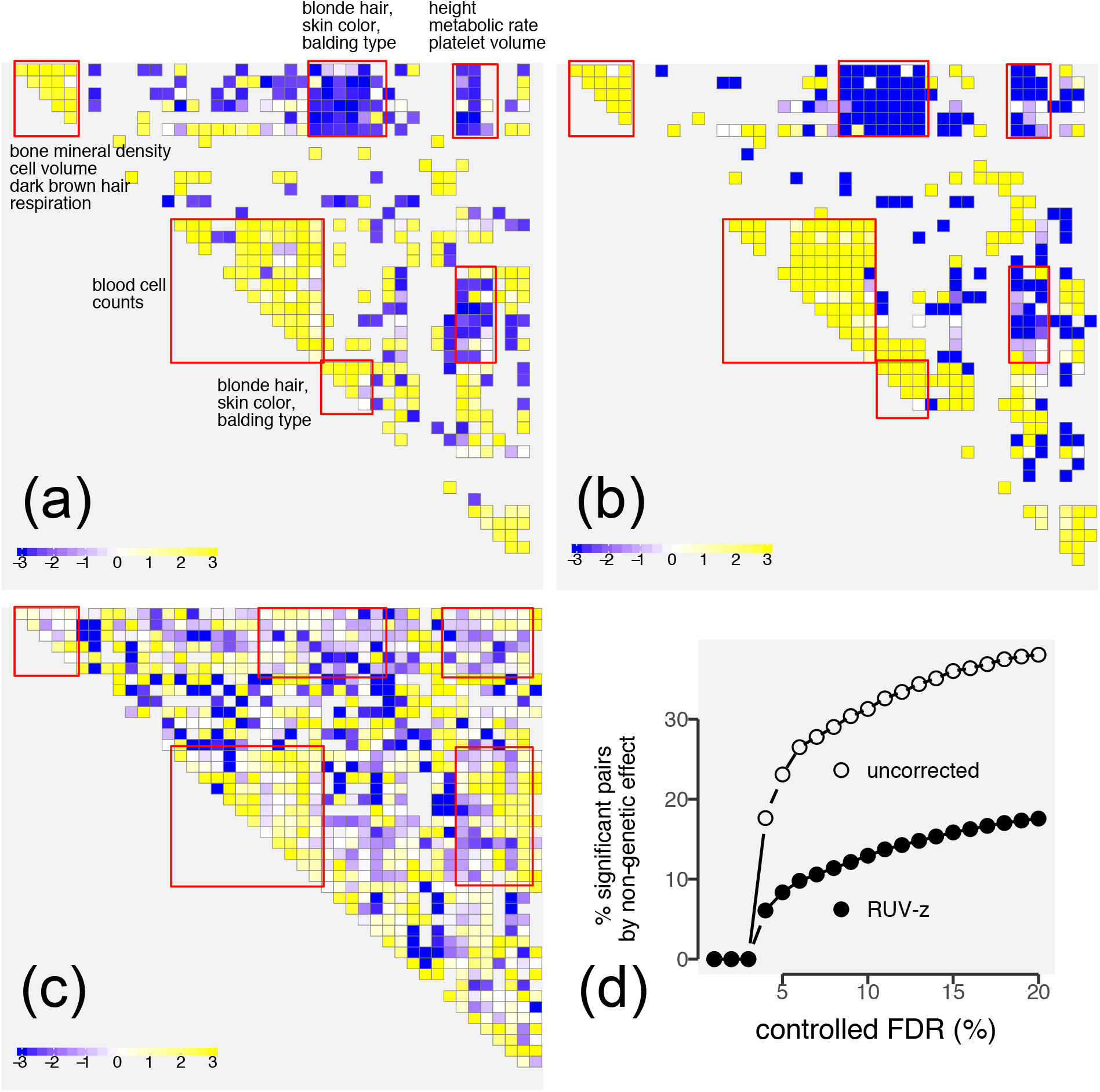
Average covariance structure between traits inferred from 47 UK biobank GWAS statistics. The averages were taken over significantly correlated pairs (at FDR < 5%) across all the LD blocks. **(a)** average covariance structure between traits using z-score based calculation without any adjustment (Eq.6); **(b)** average covariance structure of the non-genetic components (Eq.6) identified by Alg.1; **(c)** average of sparse multivariate covariance structure (Eq.17), excluding the contributions from the non-genetic covariates. **(d)** The uncorrected genetic covariance may involves higher fraction of false discoveries due to non-genetically confounded effect. We show fractions of putative false discovery pairs due to non-genetic confounders at different levels of the FDR values.

Second, since correlation structures of non-genetic confounders should occur repeatedly across many LD blocks, the directionality is expected to be consistent as well. Confounding effects of the UKBB traits consistently act in the same direction (Fig.5b), and as consequence, we also observe a similar level of consistency across many LD blocks in the marginal covariance matrix (Fig.5a). On the other hand, the genetic correlation of the sparse effects frequently flip directionality, leading to weaker correlation on average (Fig.5c).

We further reassure in the results that the fraction of potential false discovery pairs (strong correlation by the non-genetic confounders) can be more then twice higher unless we remove non-genetic sharing from the z-score matrix (Fig.5d). We empirically calibrated FDR of the covariance z-scores using the `fdrtool` package implemented in `R` [40]. And we also acknowledge that it is still possible to have trait-trait correlations may derive from both genetic, non-genetic effects, and even gene-environment interactions.

#### d. A large proportion of local pleiotropic effects are balanced across LD blocks

From this analysis, we gain novel insights into pleiotropy. We found pleiotropic relationships are frequently balanced between positive and negative interactions (Fig.6), e.g., the pairs between blood-related traits. On average across genome, such pleiotropic patterns may appear less salient than the unbalanced, therefore unidirectional, pleiotropic pairs such as the one between hair color and skin color. Unidirectional pleiotropy across genomic regions may become detectable in global pleiotropy analysis (e.g., [6]), but perhaps, the net effect of the balanced pleiotropy may not be obviously observable in comorbidity networks [18, 26, 33].

**FIG. 6:**
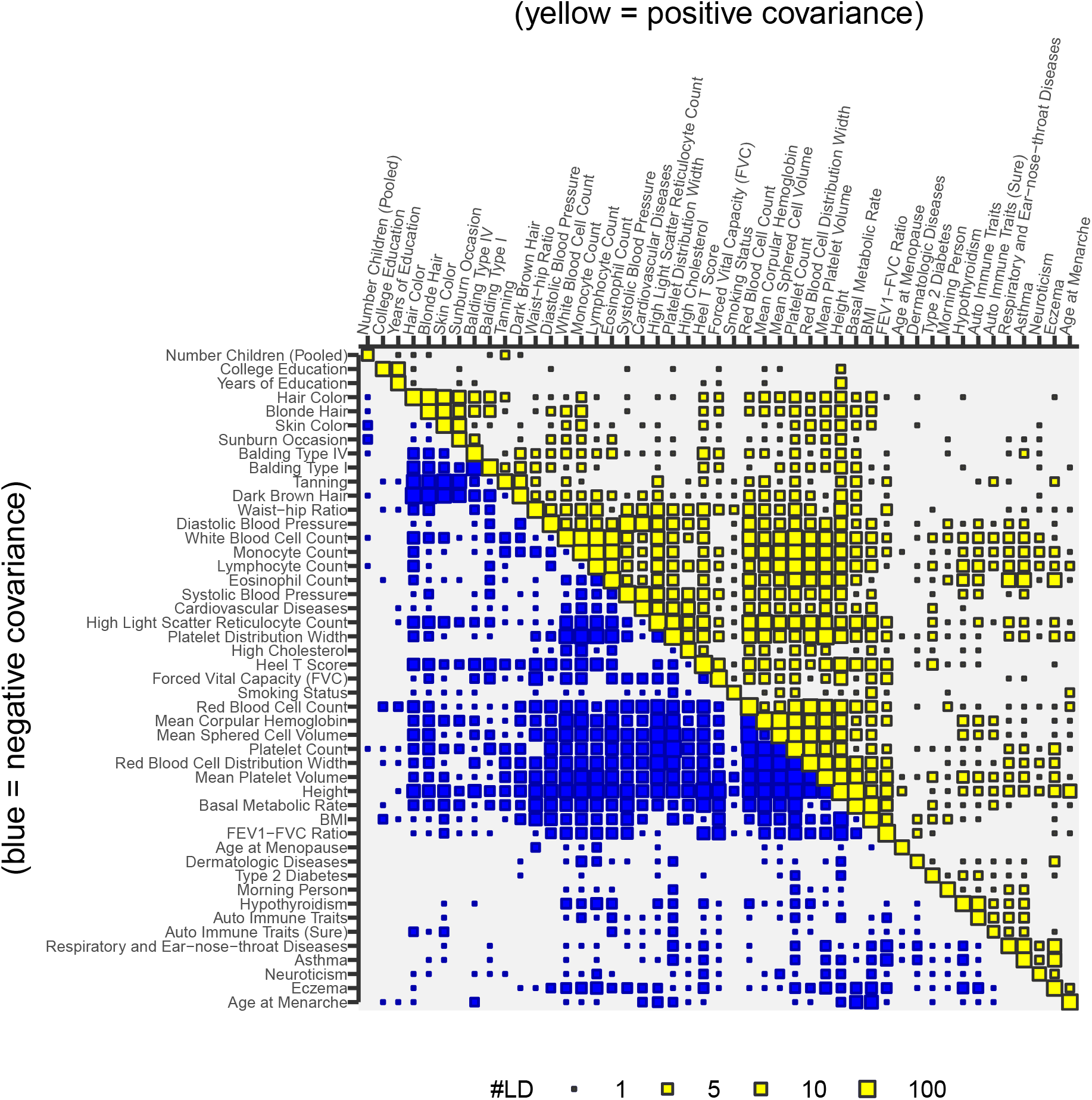
A frequency map of significant trait-trait covariance pairs.

#### e. Sparse covariance analysis recapitulates known pleiotropy and disease comorbidity

This UKBB data may not be ideal to investigate pleiotropy, as they only represents a small and biased subset of overall landscape of the biobank traits; nevertheless, the inferred pleiotropic frequency map recapitulates well-accepted comorbidity groups. For a better illustration, we visualize a subset of traits separately (Fig.7) and summarize several groups of pleiotropic modules in the following list.

(1) Common immune disorders and allergy: We confirm positive pleiotropy among eczma, neuroticism score, asthma and other respiratory problems. There are some overlap in the cases of asthma and respiratory diseases.
(2) Auto-immune diseases: Positive relationships within auto-immune disorders and hypothy-rodism, and between autoimmune traits and asthma / respiratory problems; auto-immunity is a trigger mechanism of hypothyrodism [8].
(3) Age at Menarche and BMI (body mass index): Women with earlier age of menarche tend to exhibit higher BMI, possibly due to lower metabolic rate, or vice versa; this relationship is corroborated by many studies across diverse populations [1, 5, 20, 28, 29].
(4) Cardiovascular diseases (CVD) and high cholesterol level: High cholesterol level is positive correlated with the CVD risk; common causal genes, such as *PCSK9*, have been identified by population-level genetic studies [21, 22]; positive correlation with systolic / diastolic blood pressure is somewhat obvious, but correlation with platelet distribution width had not been considered until recently [9].
(5) Pleiotropy between auto-immune diseases and immune cell types: Lymphocyte and monocyte counts are positively correlated with the immune and auto-immune disorders, stronger than other blood cell traits.

**FIG. 7:**
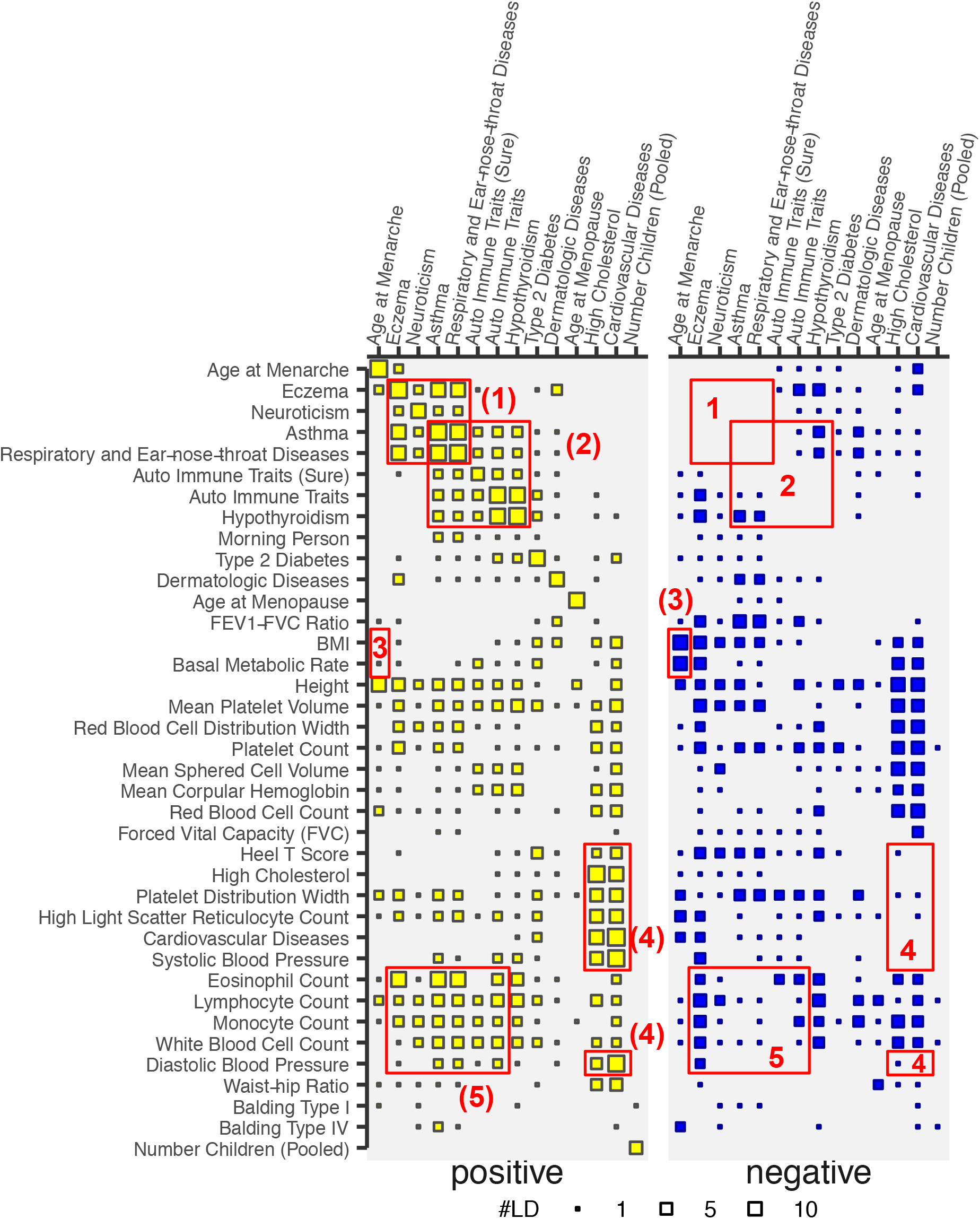
A subset of the frequency matrix of significant trait-trait covariance pairs (Fig.6), highlighting interacting partners of the disease and reproduction traits (FDR < 5%). **(1)** Common immune disorders and allergy; **(2)** Auto-immune diseases; **(3)** Age at Menarche and BMI; **(4)** Cardiovascular diseases and high cholesterol level; **(5)** Pleiotropy between auto-immune diseases and immune cell types. See the text for the brief descriptions.

## V. DISCUSSION

GWAS summary statistics data have become one of the most popular format in exchanging the results of large-scale genetics studies [35]. Unlike sharing sensitive individual-level data through secured protocols, we may avoid difficult privacy and security issues without loosing much of information. However, it is often blindly assumed that summary statistics vectors preserve desired theoretical properties, rejecting any possibilities of confounding effects.

Here, we present a novel Bayesian framework, in which a researcher can execute ML routines on GWAS summary statistics. Using this method, we particularly address two types of prevalent confounding effects-polygenic bias and non-genetic confounders. We demonstrated unless they are properly removed, test statistics of genetic variance and covariance estimation can be dramatically shifted both in simulations and real-world applications. It is not difficult to imagine that locally present minuscule bias can be easily accumulated over a thousands of LD blocks.

Our confounder correction strategy is largely inspired by existing methods in genomics [11, 36] and astrophysics [37]. In genomics, the RUV methods (removing unwanted variance) identify “control” genes or samples, which are not affected by case-control labels, and perform principal component analysis (PCA) on the control data to ascertain biases introduced by technical covariates. The half-sibling regression [37] also adopts a similar idea, but without PCA, measurements of control variates are directly included in a regression model.

Sparse modeling of genetic effects proves to be indispensable in high-dimensional multivariate analysis. We use an element-wise spike-slab prior [27] to handle SNP-level sparsity; a group-wise spike-slab prior [15] to select relevant ranks in matrix factorization. However, our sparse modeling only provide a limited resolution of “fine-mapping” of causal variants (see the supplementary for details). Improving upon current variational inference with fully factored surrogate distributions [7], we may consider more suitable variational approximations [44] in the future version of our packages.

## Acknowledgements

The authors thank Abhishek Sarkar for suggesting UK biobank data generated by BOLT-LMM and helpful discussions.

